# Competency assessment of the medical interns and nurses and prevailing practices to provide family planning services in teaching hospitals in three states of India

**DOI:** 10.1101/517326

**Authors:** Madhu Gupta, Madhur Verma, Kiranjit Kaur, Kirti Iyengar, Tarundeep Singh, Anju Singh

**Affiliations:** Department of Community Medicine and School of Public Health, Postgraduate Institute of Medical Education and Research, Chandigarh, India; United Nations Population Fund, New Delhi, India; Department of Obstetrics and Gynaecology, Postgraduate Institute of Medical Education and Research, Chandigarh, 160012. India; Department of Community Medicine, Kalpana Chawla Government Medical College, Karnal, Haryana, India

**Keywords:** contraception, competency, family planning, medical interns, nurses

## Abstract

**Objectives:** The objectives of the study was to assess the knowledge and skills of medical interns and nurses regarding family planning (FP) services, and document the prevailing FP practices in the teaching hospitals in India.

**Study Design:** A cross-sectional study was conducted in three states (Delhi, Rajasthan, and Maharashtra) of India, among randomly selected 163 participants, including medical interns (n=81) and in-service nurses (n=82), during 2017. Semi-structured, pre-tested interview schedule, was used to assess the knowledge and status of training received; and objective structured clinical examination (OSCE) based checklist was used to assess the skills.

**Results:** About 60% of the interns and 48% of the nurses knew more than five contraceptives that could be offered to the clients. About 22% (11.1% interns and 33.3% nurses) respondents believed that contraceptives should not be given to a married woman coming alone, and 31.9% (17.3% interns and 46.3% nurses) respondents reported that it was illegal to provide contraceptives to unmarried people. Nearly 43.3% interns and 69.5% nurses refused to demonstrate intrauterine contraceptive device (IUCD) insertion in the dummy uterus as per OSCE, and among those who did, 12.3% interns and 18.3% nurses had failed. About 63% interns and 63.4% of nurses had observed IUCD insertion, and 12.3% interns and 17.1% had performed IUCD insertion, during their training.

**Conclusions:** Knowledge and skills of interns and nurses regarding FP services were only partial. The medical training during graduation or internship, and during the job, was found to be inadequate to provide quality FP services.

**Implications:** The partial knowledge and skills of medical interns and nurses regarding family planning services indicated inadequate training received, and substandard quality of services rendered by them, which may put the universal access to sexual and reproductive health care services and rights in the developing countries at risk.

## 1. Introduction

A comprehensive sexual and reproductive health deals with issues concerning the reproductive system, including sexual health and not only birth control [i]. It is essential for ensuring universal access to sexual and reproductive health care services and rights, to reduce maternal mortality ratio to less than 70 per 100,000 live births and to end preventable deaths of new-borns and under-five children and for the achievement of Sustainable Development Goal three by 2030 [ii]. Despite the fact that India was the first country to launch family planning program in the year 1952, it continues to be the second most populous country in the world[iii]. In the last decade, there were minimal improvements in the family planning (FP) indicators. Data from the national level surveys suggest that the birth rate has declined from 23.8/1000 mid-year population in 2005 to 20.4/1000 mid-year population in 2016, total fertility rate from 2.7 to 2.2, contraception use rate has increased from 56.3% to 57.2%, and unmet need of FP decreased from 13.9 to 12.9 [iv,v]. There is evidence that the cafeteria or basket of choices approach of providing FP services, is not being implemented effectively [vi,vii].

To deliver effective FP services there is an urgent need to develop trained human resource that includes doctors and nurses. As per the Medical Council of India, it is necessary for the medical graduates to undergo a compulsory one-year rotatory internship after the final year to acquire the essential skills. Similarly, Indian Nursing Council ensures uniform standards of training for nurses including general midwifery and nursing (GNM), auxiliary midwifery and nursing (ANM) and Bachelor of Science in Nursing (BSc). However, it has been observed that teaching practices in medical and nursing colleges in India are often not evidence-based, and may not align with ever evolving standard protocols and the national guidelines, especially with respect to FP services [viii,ix]. With this background, this study was conducted with the objectives to assess the knowledge and skills of medical interns and nurses regarding FP services; and to document the prevailing FP practices in the teaching medical facilities in India, so that medical education could be strengthened and family planning services were improved.

## 2. Methodology

A cross-sectional study was conducted in three purposively selected states including Delhi, Rajasthan, and Maharashtra, India, between November and December 2017. The study participants were medical interns who had passed final exam of bachelor of medicine and surgery (MBBS), and completed compulsory rotatory training in the department of obstetrics and gynecology; nurses, who were already in the job, with less than 5 years’ of experience in the medical college or health posts attached with the medical colleges; and faculty or medical officer-in-charge of FP centre in the department of obstetrics and gynaecology or community medicine.

Assuming knowledge percentage score of medical interns and nurses regarding FP services as 50%, absolute precision as 10% and a design effect of 1.5, the sample size was calculated to be 145 using OpenEpi, version 3, open source calculator [x]. Assuming a 10% non-response rate, the final sample size is estimated to be 160 (80 interns and 80 nurses). Multistage simple random sampling technique was used to first select two districts within each study states, and then two medical colleges from the selected districts in the second stage. Private medical colleges were excluded, as it was difficult to get permission from these colleges. Since, Delhi did not have districts, hence, two medical colleges were selected randomly within it. However, the approval to conduct the study in one of the medical colleges could not be obtained, hence, the additional medical college from Maharashtra was selected. Hence, a total of six medical colleges (one medical college in Delhi, two in Rajasthan and three in Maharashtra) were included in the study. Interns and nurses who had fulfilled the inclusion criteria were enlisted and proportionate number (19, 31 and 31 interns from Delhi, Rajasthan and Maharashtra) were randomly selected. Permission to interview nurses could not be obtained from the selected medical college in Delhi. Hence, 32 nurses from Rajasthan and 50 from Maharashtra were randomly selected. After ensuring anonymity, written informed consent was obtained from the participants prior to the start of the interview.

A semi-structured, pre-tested interview schedule was developed in consultation with family planning experts from UNFPA, experts from the department of obstetrics and gynecology, community medicine and public health, to collect data by face to face interview. The content validity was further established by the another group of experts from the same departments. Translation of the tool was done in Hindi to interview nurses. To assess the skills of the interns and nurses an observation checklist based upon Objective Structured Clinical Examination (OSCE) was developed, for demonstration of steps of the intrauterine contraceptive device (e.g., CuT) insertion in a model, condom use on the thumb and use of Medical Eligibility Criteria (MEC) Wheel. A tool to assess the prevailing FP practices in the department of obstetrics and gynecology or community medicine was also prepared. Two postgraduate doctors (Doctor of Medicine in Community Medicine and Masters in Public Health) were specially recruited, and trained by the experts (faculty) from obstetrics and gynecology, and community medicine department to collect the data after obtaining written consent. Their work was regularly supervised by these experts in the study states to ensure data quality. Data was entered and analyzed in Statistical Package for Social Sciences, version 16.0.

### 2.1 Ethical Considerations

Institute’s Ethics Committee of the main coordinating institute and study medical colleges had approved the study.

#### Results

A total of 81 interns and 82 nurses were enrolled in the study from Rajasthan (31 interns, 32 nurses), Maharashtra (31 interns, 50 nurses) and Delhi (19 interns). Mean (standard deviation) age of interns were 23.8 (±1.2), and nurses 29.2 (±1.2)years. Males (50.6%) and females (49.3%) were equally represented among interns, while females (66.8%) were more among nurses. **(Table 1).**

**Table 1.** Background characteristics of study participants.

| **Characteristics** | **Interns**<br>**n=81 (%)** | **Nurses**<br>**n=82 (%)** | **Total**<br>**n=163 (%)** | **P value** |
| --- | --- | --- | --- | --- |
| **Sex** |  |  |  |  |
| Male | 41 (50.6) | 13 (15.8) | 54 (33.1) | 0.593 |
| Female | 40 (49.3) | 69 (84.1) | 109 (66.8) |  |
| **Mean Age (standard deviation)** | 23.8(1.2) | 29.2(3.3) | 26.4(3.7) |  |
| **Age group** |  |  |  |  |
| 20-24 | 66 (81.5) | 4 (4.9) | 70 (42.9) | 0.000 |
| 24-29 | 15(18.5) | 45 (54.9) | 60(36.8) |  |
| 30-34 | 0 | 30 (36.6) | 30(18.4) |  |
| >=35 | 0 | 3 (3.6) | 3(1.8) |  |
| **Years of experience** |  |  |  |  |
| <=1 year | 81 (100) | 2 (2.4) | 83(50.9) | 0.000 |
| 2 years | 0 | 22 (26.8) | 22(13.5) |  |
| 3 years | 0 | 16 (19.5) | 16(9.8) |  |
| 4 years | 0 | 24 (29.3) | 24(14.7) |  |
| >=5 years | 0 | 18 (22.0) | 18(11.0) |  |
| **Marital status** |  |  |  |  |
| Married | 7 (8.6) | 70 (85.4) | 77 (47.2) | 0.000 |
| Unmarried | 74 (91.4) | 12 (14.6) | 86 (52.8) |  |
| **State** |  |  |  |  |
| **Delhi** |  |  |  |  |
| * MC1 | 19 (23.5) | 0 | 19 (11.7) |  |
| **Rajasthan** |  |  |  |  |
| MC 2 | 16 (19.8) | 16 (19.8) | 32 (19.6) |  |
| MC 3 | 15 (18.5) | 16 (19.8) | 31 (19.0) |  |
| **Maharashtra** |  |  |  |  |
| MC 4 | 11 (13.6) | 19 (23.5) | 30 (18.4) |  |
| MC 5 | 17 (21.0) | 23 (28.4) | 40 (24.5) |  |
| MC 6 | 3 (3.7) | 8(9.9) | 11 (6.7) |  |

\*MC : Medical College

### 2.1. Knowledge of interns and nurses regarding contraceptive methods

About 60% of the interns and 48% of the nurses knew about more than 5 contraceptives that could be offered through the cafeteria approach. Majority of the respondents were of the opinion that condoms (88.3%) and oral contraceptive pills (OCPs) [77.9%] were the best contraceptives for newly married couples. **(Table 2).** For a woman with one child, intrauterine contraceptive device (IUCD) [93.4%], and for a woman with three children, sterilization (95.8%) was the most common response. About one-fifth of the participants (22%) responded that the contraceptives should not be given to a married woman who was coming alone. Nearly 31.9% (17.3% interns and 46.3% nurses) respondents told that it was illegal to provide the contraceptives to unmarried people. Knowledge of interns and nurses regarding oral contraceptive pills, condoms, and emergency contraceptives is presented in **table 3.** Respondents knew the common medical conditions like cardiovascular diseases (41.7%), breast diseases (30%), headache/migraine (25.7%) and thromboembolic disorders (22.1%) to rule out before prescribing OCPs. Knowledge of interns and nurses regarding reversible long-acting contraceptives (IUCDs and hormonal contraceptives) and permanent contraceptives (tubectomy) is shown in **table 4.** Duration of protection (10 years) offered by CuT 380 A was known to 51.9% interns and 35.4% nurses. About 38% were aware that depot medroxyprogesterone acetate (DMPA) is available in the government supply. About 83% of respondents had seen the eligibility checklist for tubectomy during their training period, but only half of them (43%) had seen tubectomy operation. Similarly, the fact that amenorrhea, exclusive breastfeeding and six months of the postpartum period are the three essential prerequisites for lactational amenorrhea (LAM) to be an effective contraceptive method, was known to 80%, 63.8%, and 48.5% respondents, respectively. Overall, the assessment of knowledge in terms of correct, partial and wrong is presented in **Table 5.**

**Table 2.**
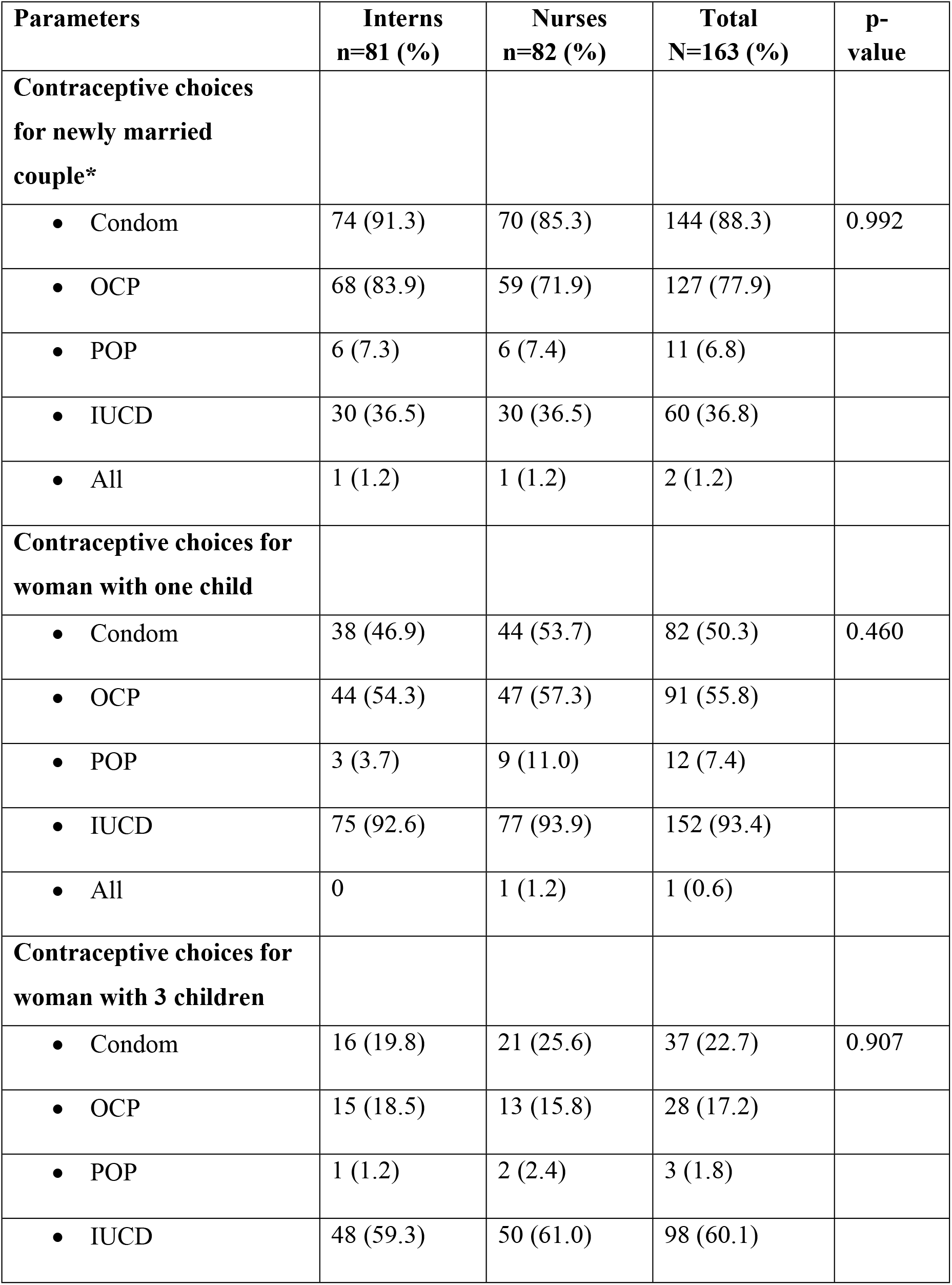
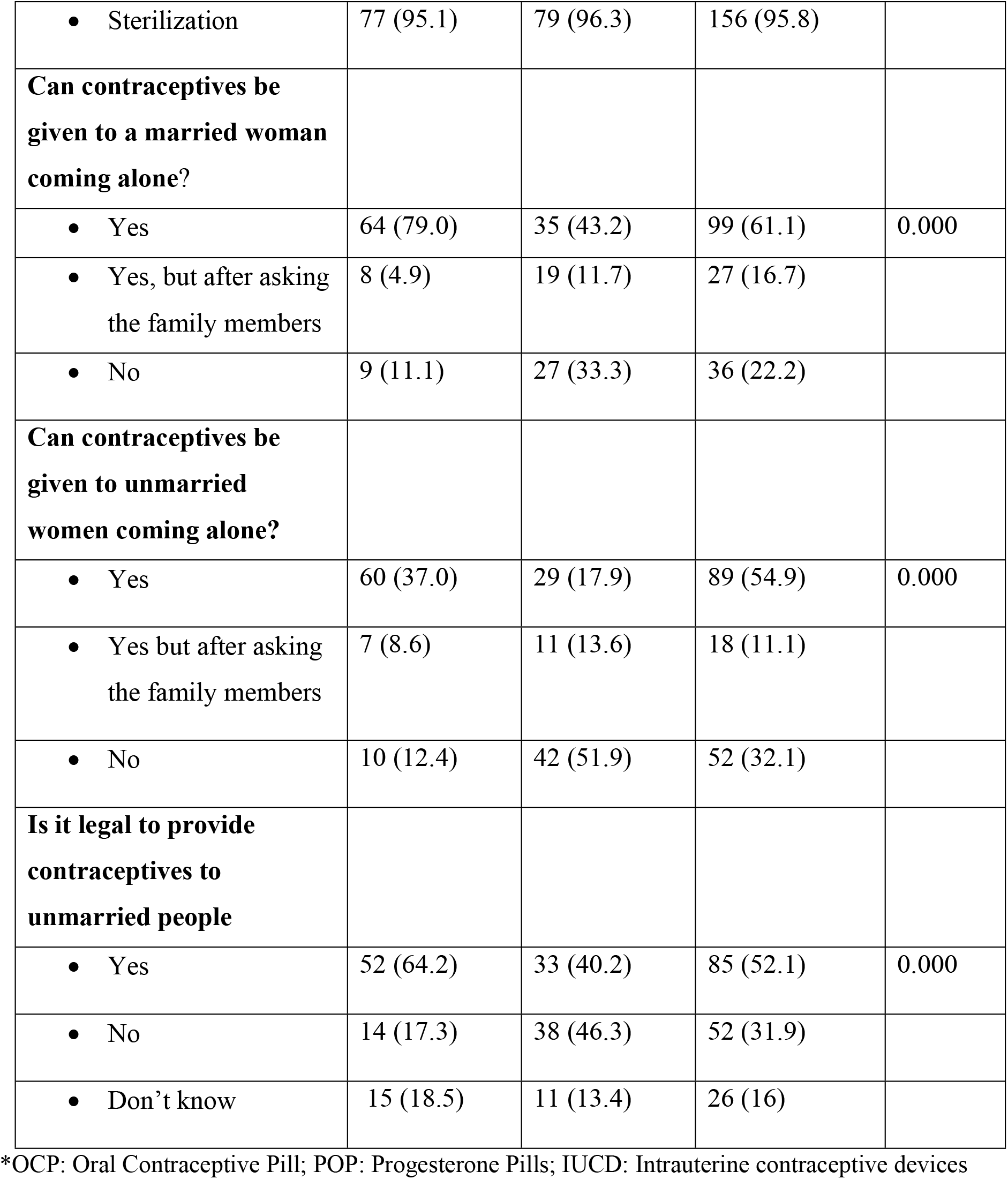
Knowledge of interns and nurses regarding various contraceptive methods and reproductive rights of the clients.

**Table 3.** Knowledge of interns and nurses regarding oral contraceptive pills, condoms and emergency contraceptives.

| **Contraceptives** | **Interns**<br>**n=81 (%)** | **Nurses**<br>**n=82 (%)** | **Total**<br>**N=163 (%)** | **p-**<br>**value** |
| --- | --- | --- | --- | --- |
| **Oral contraceptives Pills**<br>**(OCPs)** |  |  |  |  |
| *Conditions to rule out from*<br>*history before prescribing*<br>*OCPs* |  |  |  |  |
| • Smoking | 10 (12.4) | 5 (6.1) | 15 (9.2) | 0.060 |
| • Diabetes | 10 (12.4) | 15 (18.3) | 25 (15.3) |  |
| • Headaches | 24 (29.6) | 18 (21.95) | 42 (25.7) |  |
| • Cardiovascular diseases | 45 (55.6) | 23 (28.1) | 68 (41.7) |  |
| • Thromboembolic disorders | 29 (35.8) | 7 (8.5) | 36 (22.1) |  |
| • Post-partum haemorrhage | 7 (8.6) | 2 (2.4) | 9 (5.5) |  |
| • liver disease | 28 (34.6) | 11 (13.4) | 39 (24.0) |  |
| • Breast disease | 33 (40.7) | 16 (19.8) | 49 (30.6) |  |
| *Instructions to be given while*<br>*prescribing OCPs should*<br>*include* |  |  |  |  |
| • When to start OCP | 64 (79.0) | 62 (76.6) | 126 (77.8) | 0.471 |
| • Daily intake without fail | 71 (87.7) | 67 (82.7) | 138 (85.2) |  |
| • What to do if misses a pill | 56 (69.5) | 41 (50.6) | 97 (59.9) |  |
| • Possible side effects | 22 (27.5) | 13 (16.1) | 35 (21.6) |  |
| ***What to do if client misses 2 pills?*** |  |  |  |  |
| To take 2 pills the next day | 42 (51.9) | 43 (53.1) | 85 (52.8) | 0.025 |
| Again 2 pills the second next day | 16 (19.8) | 12 (14.9) | 28 (17.3) |  |
| The couple should also use condom for 7 days | 34 (42.0) | 12 (14.9) | 46 (24.4) |  |
| ***Can OCPs be given to (yes)*** |  |  |  |  |
| Newly married women | 68 (84.0) | 43 (52.5) | 111 (68.1) | 0.079 |
| Illiterate women | 64 (79.0) | 60 (73.2) | 124 (76.1) |  |
| Women who do not want any more children | 53 (65.4) | 61 (74.4) | 114 (70.0) |  |
| **Condoms** |  |  |  |  |
| ***Failure rate of Condoms if used correctly*** |  |  |  |  |
| <5% | 34 (2.0) | 17 (21.0) | 51 (31.5) | 0.002 |
| 6-15% | 35 (43.2) | 20 (24.7) | 55 (34.0) |  |
| >15% | 4 (5.0) | 5 (6.2) | 9 (5.6) |  |
| Other | 1 (1.2) | 6 (7.4) | 7 (4.3) |  |
| Do not know | 3 (3.7) | 12 (14.8) | 15 (9.3) |  |
| ***Two most common Advantages of condom*** |  |  |  |  |
| Dual protection against pregnancy and STI/HIV<sup>^</sup> | 70 (86.4) | 47 (58.0) | 117 (72.2) | 0.016 |
| • No side effects | 23 (28.4) | 34 (42.0) | 57 (35.2) |  |
| **Emergency contraceptives** |  |  |  |  |
| *Type of contraception used after unprotected intercourse* |  |  |  |  |
| • Emergency contraceptive pills | 76 (93.8) | 71 (86.6) | 147 (90.2) | 0.000 |
| • Intrauterine contraceptive devices | 45 (55.6) | 6 (7.3) | 51 (31.3) |  |
| • Yuzpe's method | 24 (29.6) | 4 (4.9) | 28 (17.2) |  |
| *Emergency contraceptive pill is effective if consumed within* |  |  |  |  |
| • 24 hours of unprotected intercourse | 0 | 2 (2.4) | 2 (1.2) | 0.177 |
| • 48 hours of unprotected intercourse | 3 (3.7) | 6 (7.32) | 9 (5.5) |  |
| • 72 hours of unprotected intercourse | 77 (95.1) | 68 (82.9) | 145 (89.0) |  |
| **Frequency for taking centchroman** |  |  |  |  |
| • 1 tablet weekly | 18 (22.2) | 2 (2.4) | 20 (12.3) | 0.000 |
| • Do not know | 48 (59.3) | 80 (97.6) | 128 (78.5) |  |

^STI/HIV: Sexually Transmitted Infections/Human Immunodeficiency Virus

**Table 4.** Knowledge of interns and nurses regarding reversible long acting contraceptives (intrauterine contraceptive devices and hormonal contraceptives, post-partum contraception) and permanent contraceptives (tubectomy).

| **Parameters** | **Interns**<br>**n=81 (%)** | **Nurses**<br>**n=82**<br>**(%)** | **Total**<br>**n=163 (%)** | **P**<br>**value** |
| --- | --- | --- | --- | --- |
| **Awareness about types of intrauterine contraceptive devices (IUCDs)** |  |  |  |  |
| • Copper containing IUCDs | 81 (100) | 78 (95.1) | 159 (97.6) | 0.000 |
| • Hormonal IUCDs | 60 (74.1) | 12 (14.6) | 72 (44.2) |  |
| • First generation /inert IUCDs | 51 (63.0) | 23 (28.1) | 74 (45.4) |  |
| **Duration of protection offered by CuT 380 A** |  |  |  |  |
| • 10 Years | 42 (51.9) | 29 (35.4) | 71 (43.6) | 0.034 |
| **Most common conditions to rule out before inserting Copper T** |  |  |  |  |
| • Pregnancy | 46 (56.8) | 40 (48.8) | 86 (52.8) | 0.287 |
| • STI/HIV | 53 (65.4) | 36 (43.9) | 89 (54.6) |  |
| • Irregular Periods | 50 (61.7) | 54 (65.9) | 104 (63.8) |  |
| • Adnexal Mass/Ectopic Pregnancy | 41(50.6) | 25 (30.5) | 66 (40.5) |  |
| • Multiple Sexual Partners | 1 (1.2) | 0 | 1 (0.6) |  |
| **Most common side effects of Cu-T insertion** |  |  |  |  |
| • Pain/cramps | 61 (75.3) | 55 (67.9) | 116 (71.6) | 0.004 |
| • Bleeding/menorrhagia/spotting/irregular bleeding | 61 (75.3) | 72 (88.9) | 133 (82.1) |  |
| • Infections/PID/vaginal discharge | 51 (63.0) | 32 (39.5) | 83 (51.2) |  |
| • Expulsion | 29 (34.6) | 9 (11.1) | 37 (22.8) |  |
| **When is Post-Partum IUCD (PPIUCD) to be inserted** |  |  |  |  |
| • Within 10 minutes of delivery | 37 (45.7) | 30 (36.4) | 67 (41.1) | 0.001 |
| • Within 48 hours | 27 (33.3) | 10 (12.2) | 37 (22.7) |  |
| • During caesarean section | 10 (12.4) | 8 (9.8) | 18 (11.0) |  |
| • Other | 26 (32.1) | 11 (13.4) | 37 (22.7) |  |
| • Don't know | 2 (2.5) | 13 (16.0) | 15 (9.3) |  |
| **Time for taking consent for PPIUCD insertion** |  |  |  |  |
| • Ante-natal period | 40 (49.4) | 29 (35.8) | 69 (42.3) | 0.001 |
| • Intra-natal period | 31 (38.3) | 16 (19.5) | 45 (28.8) |  |
| • Post-natal period | 8 (9.9) | 25 (30.9) | 33 (20.4) |  |
| **Questions to ask before inserting Depot Medroxy Progesterone Acetate (DMPA)** |  |  |  |  |
| • Pregnancy | 14 (17.3) | 12 (14.8) | 26 (16.0) | 0.639 |
| • Irregular periods | 24 (29.6) | 15 (18.5) | 39 (24.7) |  |
| • Breast cancer | 20 (24.7) | 12 (14.9) | 32 (19.8) |  |
| • Liver diseases | 9 (11.1) | 5 (6.2) | 14 (8.6) |  |
| • Thromboembolic episodes | 11 (13.6) | 7 (8.6) | 18 (11.1) |  |
| **Topics to be covered while counselling for DMPA** |  |  |  |  |
| • Menstruation related side effects | 27 (33.3) | 14 (17.2) | 41 (25.2) | 0.668 |
| • Delayed return of fertility | 14 (17.3) | 5 (6.1) | 19 (11.6) |  |
| • Don't know | 19 (23.7) | 12 (15.0) | 31 (19.4) |  |
| **Contraceptives that can be advised to breast feeding mother** |  |  |  |  |
| • Intrauterine contraceptive device | 45 (55.6) | 61 (74.4) | 106 (65.0) | 0.000 |
| • Injectable contraceptives | 8 (9.9) | 9 (11.0) | 17 (10.4) |  |
| • Progesterone only Pills | 31 (38.3) | 6 (7.3) | 37 (22.7) |  |
| • Condoms | 56 (69.1) | 54 (65.9) | 110 (67.9) |  |
| **Tubectomy: Have ever seen (yes)** |  |  |  |  |
| • Eligibility checklist for Tubectomy | 12 (85.2) | 14 (79.3) | 26 (82.2) |  |
| • Tubectomy operation | 31 (38.3) | 39 (47.6) | 70 (43.0) |  |
| • Consent form for Tubectomy | 24 (29.6) | 50 (61.0) | 74 (45.4) |  |
| **Prerequisites for lactational amenorrhea to be an effective contraceptive** |  |  |  |  |
| • Amenorrhoea | 18 (22.2) | 14 (83.0) | 32 (80.4) | 0.024 |
| • Exclusive breast feeding | 59 (72.8) | 45 (54.9) | 104 (63.8) |  |
| • 6 months post-partum | 43 (53.1) | 36 (43.9) | 79 (48.5) |  |
| • Don't know | 11 (13.8) | 27 (33.3) | 38 (23.6) |  |

**Table 5.** Overall knowledge of interns and nurses regarding various contraceptive methods

| Parameters* | Response of Interns n=81 (%) |  |  |  | Response of Nurses n=82 (%) |  |  |  | p value |
| --- | --- | --- | --- | --- | --- | --- | --- | --- | --- |
|  | Correct | Partially correct | Wrong | Don't know | Correct | Partially correct | Wrong | Don't know |  |
| Contraceptive choices for newly married couple (Condom, OCP, POP, IUCD, all, few of these) | 23<br>(28.3) | 50<br>(61.7) | 8<br>(9.8) | 0 | 23<br>(28.0) | 36<br>(43.0) | 31<br>(37.8) | 0 (0) | 0.000 |
| Contraceptive choices for woman with 1 child (Condom, OCP, POP, IUCD, all, few of these) | 27<br>(33.3) | 28<br>(34.5) | 26<br>(32.1) | 0 | 35<br>(42.3) | 23<br>(28) | 23<br>(28.0) | 1(1.2) | 0.440 |
| Contraceptive choices for woman with 3 children (Condom, OCP, POP, IUCD, sterilization, all, few of these) | 22<br>(27.1) | 27<br>(33.3) | 31<br>(38.2) | 1<br>(1.2) | 26<br>(31.7) | 27<br>(32.1) | 29<br>(34.1) | 0 (0) | 0.707 |
| Can contraceptives be given to a married woman coming alone? | 68<br>(83.9) | 0 | 13<br>(16.0) |  | 37<br>(45.1) | 0 | 45<br>(54.8) | 0 (0) | 0.000 |
| Can Contraceptives be given to unmarried women coming alone? | 61<br>(75.3) | 0 | 18<br>(22.2) | 2<br>(2.4) | 32 (39) | 0 | 48<br>(58.5) | 2 (2.4) | 0.000 |
| Is it legal to provide contraceptives to unmarried people? | 52<br>(64.1) | 0 | 14<br>(17.2) | 15<br>(18.5) | 33<br>(40.2) | 0 | 38<br>(46.3) | 11<br>(13.4) | 0.000 |
| Types of IUDs available (1<sup>st</sup> generation, Copper, Hormonal) | 18<br>(22.2) | 15<br>(18.5) | 48<br>(59.3) | 0 | 51<br>(62.2) | 17<br>(20.7) | 9<br>(11.0) | 1 (1.2) | 0.000 |
| Common contraindications for IUDs (pregnancy, STI/HIV, irregular periods/ adnexal mass, ectopic | 17<br>(21.0) | 18<br>(22.0) | 42<br>(51.9) | 3<br>(3.7) | 20<br>(24.4) | 27<br>(32.9) | 25<br>(30.5) | 8 (9.8) | 0.035 |
| pregnancy, multiple sexual partners) |  |  |  |  |  |  |  |  |  |
| Common side effects of IUDs (cramps, menstrual problems, infections, RTI/STI, expulsion) | 12<br>(14.8) | 31<br>(38.3) | 36<br>(44.1) | 1<br>(1.2) | 11<br>(13.4) | 38<br>(46.3) | 26<br>(32.7) | 5 (6.1) | 0.16<br>9 |
| Types of Cu-T in government supply (CuT375, 380A) | 24<br>(29.6) | 48<br>(59.3) | 3<br>(3.7) | 6<br>(7.4) | 19<br>(23.2) | 28<br>(34.1) | 6 (7.3) | 29<br>(35.4) | 0.00<br>0 |
| How long does Cu-T 380A offer protection? | 42<br>(51.9<br>1) | 0 (0) | 34<br>(42.0) | 5<br>(6.2) | 29<br>(35.4) | 0 (0) | 37<br>(45.1) | 16<br>(19.5) | 0.01<br>6 |
| When is PPIUCD inserted? | 15<br>(18.5) | 43<br>(53.1) | 11<br>(13.6) | 12<br>(14.8) | 9<br>(11.0) | 36<br>(43.9) | 8 (9.8) | 29<br>(35.4) | 0.02<br>2 |
| When is consent for PPIUCD taken? | 61<br>(75.3) | 0 (0) | 10<br>(12.3) | 10<br>(12.3) | 3 (3.7) | 43<br>(52.4) | 17<br>(20.7) | 19<br>(23.2) | 0.00<br>0 |
| Medical contraindications for OCPs (Any 4 correct responses) | 21<br>(25.9) | 0 (0) | 49<br>(60.5) | 11<br>(13.6) | 15<br>(17.3) | 0 (0) | 35<br>(42.7) | 32<br>(39.0) | 0.00<br>1 |
| Yes OCPs can be bought over the counter | 53<br>(65.4) | 0 (0) | 24<br>(29.6) | 4<br>(4.9) | 52<br>(62.3) | 0 (0) | 29<br>(35.4) | 1 (1.2) | 0.32<br>1 |
| Instructions given before starting OCPs | 48<br>(59.3) | 27<br>(33.3) | 4<br>(4.9) | 2<br>(2.5) | 38<br>(46.3) | 32<br>(39.0) | 5 (6.1) | 7 (8.5) | 0.21<br>5 |
| Instructions if 2 pills are missed | 10<br>(12.3) | 29<br>(35.8) | 20<br>(24.7) | 22<br>(27.2) | 4 (4.9) | 25<br>(30.5) | 22<br>(26.8) | 31<br>(37.8) | 0.21<br>4 |
| Whom can OCPs be given to? (Newly married women, illiterate women, Women who do not want any more children) | 41<br>(50.6) | 26<br>(32.1) | 14<br>(17.3) | 0 (0) | 31<br>(37.8) | 28<br>(34.1) | 22<br>(26.8) | 1 (1.2) | 0.23<br>7 |
| Which OCPs are available in govt. supply (Mala N, Mala D) | 14<br>(17.3) | 63<br>(77.8) | 3<br>(3.7) | 1<br>(1.2) | 23<br>(28.0) | 39<br>(47.6) | 5 (6.1) | 15<br>(18.3) | 0.00<br>0 |
| What kind of contraceptive is DMPA? | 59<br>(72.8) | 0(0) | 8<br>(9.9) | 14<br>(17.3) | 32<br>(39.0) | 0(0) | 3 (3.7) | 47<br>(57.3) | 0.00<br>0 |
| Medical contraindications for DMPA (Any 3 correct responses out of pregnancy/irregular periods, breast cancer, liver disease, thromboembolic episodes) | 8<br>(9.9) | 21<br>(25.9) | 11<br>(13.6) | 41<br>(50.6) | 5 (6.1) | 16<br>(19.5) | 3 (3.7) | 58<br>(70.0) | 0.03<br>1 |
| Common side effects of DMPA (menstruation related side effects, delayed return of fertility) | 9<br>(11.1) | 22<br>(27.2) | 4<br>(4.9) | 46<br>(56.8) | 4 (4.9) | 12<br>(14.6) | 2 (2.4) | 64<br>(78.0) | 0.03<br>7 |
| Yes injectable contraceptives are available in the government supply | 36<br>(44.4) | 0 | 33<br>(40.7) | 12<br>(14.8) | 28<br>(34.1) | 0 | 28<br>(34.1) | 26<br>(31.7) | 0.03<br>8 |
| Prerequisites for lactational amenorrhea (amenorrhoea, exclusive breast feeding for 6 months) | 8<br>(9.9) | 44<br>(54.3) | 15<br>(18.5) | 14<br>(17.3) | 10<br>(12.2) | 25<br>(30.5) | 17<br>(20.7) | 30<br>(36.6) | 0.01<br>0 |
| Contraceptives that can be given to breastfeeding women (Condoms, POP, IUCD, Injectable) | 14<br>(17.3) | 45<br>(55.6) | 16<br>(19.8) | 6<br>(7.4) | 14<br>(17.1) | 35<br>(42.7) | 26<br>(31.7) | 7 (8.5) | 0.29<br>6 |

\*OCP: Oral Contraceptive Device; POP: Progesterone Only Pill; IUCD: Intrauterine
Contraceptive Device; DMPA: Depot medroxy progesterone acetate, PPIUCD: Post Partum
Intrauterine Contraceptive Device; STI/HIV: Sexually Transmitted Infections/Human
Immunodeficiency Virus

There were non-significant differences in the knowledge as per the age and gender of the participants. However, it was observed that information regarding the legality of contraceptives to be given to unmarried people significantly (p=0.049) improved with age among nurses, as 65% of the nurses between 30-34 years have answered it correctly as compared to 32.7% and 28.6% among 25-29 years and 20-24 years age group, respectively. Female interns had significantly better knowledge as compared to male interns regarding the choice of contraceptives for women with one child (47.5% vs 24.4%; p=0.036), with three children (45% vs 17.1%; p=0.042); and regarding types of IUCDs (90% vs 65.9%; p=0.009).

### 2.2. Skills of nurses and interns regarding the use of contraceptive methods (Objective structured clinical examination)

About 19.8% interns and 64.6% nurses refused to demonstrate the use of MEC wheel for choosing the best contraceptives for hypothetical cases, as shown in **Table 6.** Further, 43.2% interns and 69.5% nurses refused, while 12.2 % nurses and 44.4% interns passed in demonstrating CuT insertion in dummy uterus. Correct steps of using a condom on the thumb were demonstrated by 63% interns and 40.2% nurses.

**Table 6.** Objective structure clinical examination (OSCE) score of interns and nurses.

| Parameters | Interns<br>n=81 (%) | Nurses<br>n=82<br>(%) | p-value |
| --- | --- | --- | --- |
| **Medical eligibility criteria (MEC) wheel demonstration** |  |  |  |
| • Pass | 57 (70.4) | 22 (26.8) | 0.148 |
| • Fail | 8 (9.9) | 7 (8.5) |  |
| • Refused to demonstrate appropriate usage of MEC | 16 (19.8) | 53 (64.6) |  |
| **Intrauterine contraceptive device (IUCD): Cu T** |  |  |  |
| • Pass | 36 (44.4) | 10 (12.2) | 0.000 |
| • Fail | 10 (12.3) | 15 (18.3) |  |
| • Refused to perform demonstration of Cu T insertion | 35 (43.2) | 57 (69.5) |  |
| **Steps for CuT insertion** |  |  |  |
| • Washes hands and wears gloves | 31 (38.3) | 15 (18.3) | 0.729 |
| • Insert the sterile sound with 'no touch' technique | 38 (46.9) | 12 (14.6) |  |
| • Load IUCD in its sterile package | 31 (38.2) | 7 (8.6) |  |
| • Set the blue depth gauge to the measurement of the uterus | 29 (35.8) | 9 (11.0) |  |
| • Carefully insert the loaded IUCD and release it into the uterus | 40 (49.4) | 14 (17.1) |  |
| • Take out the plunger | 31 (38.3) | 15 (18.3) |  |
| • Partially withdraw the insertion tube till the string are visible | 24 (29.6) | 9 (11.0) |  |
| • Use sterile scissors to cut the IUCD strings to 3-4 cm length in the vagina | 20 (24.7) | 11 (13.4) |  |
| **Barrier contraceptive usage (Condom)** |  |  |  |
| • Pass | 51 (63.0) | 33 (40.2) | 0.348 |
| • Fail | 15 (18.5) | 12 (14.6) |  |
| • Check the expiry date on the wrapper | 1 (1.2) | 3 (3.7) |  |
| **Steps for condom usage** |  |  |  |
| • Open the package without tearing the condom | 64 (79) | 42 (51.2) | 0.985 |
| • Do not use teeth to open the wrapper | 64 (79.0) | 41 (50.0) |  |
| • Hold the condom by the last ½ inch at the tip, making sure to squeeze out any air | 31 (38.3) | 19 (23.2) |  |
| • Put the condom on the tip of the thumb. | 61 (75.3) | 38 (46.3) |  |
| • While still pinching the tip, unroll the condom down the shaft, to the base | 33 (40.47) | 27 (32.9) |  |
| • Remove the condom by rolling it off | 51 (63.0) | 31 (37.8) |  |
| • Throw the condom into the dustbin | 39 (48.1) | 24 (29.3) |  |

### 2.3. Status of prevailing FP practices

Almost 90% of interns received training on FP methods and services, and 76.5% were also posted in FP clinics (**Table 7**). Although 63% interns and nurses have seen the insertion of IUCD, more nurses (17.1%) reported performing IUD insertion than interns (12.3%) at least once. A very small proportion of interns and nurses reported having seen implantable contraceptives (12.3%) and spermicides (17.3%). All the medical colleges reported conducting training of students for FP methods and services. The details pertaining to infrastructure and facilities are depicted in Supplementary Table 1. None of the colleges had a MEC wheel for training purposes. Adequate stock of FP methods (contraceptives, pregnancy test, CuT etc.) was available in almost all the colleges, but only 3/6 medical colleges reported to have an established FP logistic management information system, as depicted in Supplementary Table 2.

**Table 7.** Status of training of interns and nurses on family planning (FP) methods and services.

|  | **Interns**<br>**n=81 (%)** | **Nurses**<br>**n=82 (%)** | **p-value** |
| --- | --- | --- | --- |
| **Status of Training\*** |  |  |  |
| • Attended a class on FP methods during training | 73 (90.1) | 64 (78.0) | 0.035 |
| • Posted in FP clinic | 62 (76.5) | 38 (46.3) | 0 |
| • Observed counseling on different FP methods | 48 (59.3) | 57 (69.5) | 0.172 |
| • Ever seen MEC wheel | 10 (12.3) | 6 (7.3) | 0.281 |
| • Ever observed MEC wheel used for a patient | 6 (7.4) | 4 (4.9) | 0.501 |
| **Number of students who have ever\*** |  |  |  |
| • Observed IUCD insertion | 51 (63.0) | 52 (63.4) | 0.952 |
| • Performed IUCD insertion | 10 (12.3) | 14 (17.1) | 0.394 |
| • Inserted IUCD on dummy/ model | 9 (11.1) | 11 (13.4) | 0.645 |
| • Observed IUCD removal | 26 (32.1) | 42 (51.2) | 0.013 |
| • Performed IUCD removal | 16 (19.8) | 23 (28.0) | 0.215 |
| **Number of students who have ever seen** |  |  |  |
| • Condom | 78 (96.3) | 82 (100) | 0.079 |
| • Oral Contraceptive Pill | 78 (96.3) | 80 (97.6) | 0.64 |
| • Intrauterine Device | 80 (98.8) | 81 (98.8) | 0.993 |
| • Depot medroxy progesterone acetate | 32 (39.5) | 28 (34.1) | 0.478 |
| • Emergency Contraceptive Pill | 64 (79.0) | 63 (76.8) | 0.737 |
| • Centchroman | 29 (35.8) | 19 (23.2) | 0.077 |
| • Hormonal IUCD | 38 (46.9) | 9 (11.0) | 0 |
| • Implantable Contraceptives | 10 (12.3) | 13 (15.9) | 0.52 |
| • Spermicides | 14 (17.3) | 5 (6.1) | 0.026 |

\*MEC: Medical Eligibility Checklist; IUCD: Intrauterine contraceptive device

## 3. Discussion

This study highlights the partial knowledge and skills of the interns and nurses regarding family planning services in the medical colleges in Rajasthan, Maharashtra, and Delhi, India. The status of training regarding these services was also found to be inadequate. These results have any implications in terms of providing universal access to quality reproductive and sexual health to the clients, which may lead to a compromise in the reproductive rights of the population, especially of the women, as they are most often not the decision makers in opting for the FP methods.

Our study sample had proportionate representation from male (54, 33.1%) and female (109,66.9%) groups. A higher proportion of females were because the majority of the nurses were females (84.1%). Significantly higher knowledge of female participants as observed in this study is similar to Raselekoane NR (2016) study, who reported a non-serious approach of the male students towards contraception and family planning [11]. Gender differences should not be ignored during medical educations to avoid compromise with the sexual and reproductive health rights of the clients[12]. Amongst nurses, improvement in knowledge in certain topics with age may be attributed to change in marital status and hence clarity of certain concepts. However, it is recommended to have in job refresher training programs as a routine activity.

It was observed that participants did not report to provide FP methods as per cafeteria approach to an eligible woman [**Error! Bookmark not defined.**]. Most of them were unaware of the new FP methods [like DMPA (86%)] that have been introduced in India [13]. Rhythm method (87.9%), coitus interrupts (55.4%), LAM (45.2%) were preferred natural methods during counseling. Fehring RJ et al (Missouri, 1999), also observed that health care providers were likely to recommend calendar calculations and monitoring basal body temperature to their clients for managing pregnancies [14]. IUCD, condom, and sterilization were preferred contraceptives for newly married women, for women with one child and for women with three children. Jain AK et al (India, 2017), observed 15.6% of contraceptive users receiving information on all four items (IUCDs, condom, sterilization, and pill) [15]. In our study, 32.1% of participants reported that FP services should not be offered to unmarried females coming alone, and 31.9% of participants reported that providing FP services to unmarried people is illegal. Tilhaun et al study (Ethiopia, 2010) also observed a negative attitude toward providing FP methods to unmarried adolescents [16]. Sociocultural norms of Indian society contribute in having sex-related issues a taboo and hinder young people from seeking counseling regarding sexual health [17]. It has been reported earlier that that healthcare providers often impose unnecessary barriers in dispensing contraceptives, including denial of a contraceptive method on the basis of age, parity, marital status or lack of parental or spousal authorization [18, 19].

Knowledge regarding side effects and contraindications of IUCD use (68.7%) was better than other studies done in the past [20, 21]. This may be attributed to improvement in health promotion activities conducted in the last decade. Few interns (12%) had adequate knowledge about instructions that should be given to a woman who had missed up to two doses of her OCPs. These findings are in line with a study by Rutter W et al (Australia 1988) [22]. Adequate knowledge of interns and nurses regarding male condoms (72%) in this study is similar to the observations made by Simbar et al study (Iran 2005) [23].

More than 60% participants were clueless about injectable contraceptives particularly DMPA. Hogmark et al (India, 2013), reported that despite the positive attitude of medical students towards modern contraceptives, sex education and FP counseling, they still had misconceptions about modern methods of contraception in Maharashtra, which is consistent with the findings of the present study [24]. Knowledge about LAM was partial, as only 9.9% interns and 12.2% nurses (Supplementary table 1) could correctly enumerate three criteria of LAM (Table 4) to be an efficacious method [25]. Singh S et al (Delhi, 2002) observed high awareness about emergency contraception among doctors who felt that the use of emergency contraceptives can bring down the number of induced abortions [26]. Less than half of the respondents (61% nurses; 30% interns) had ever witnessed tubectomy operation, while 16% had seen eligibility checklist and 45.5 % had seen the consent form for tubectomy. This difference is probably because taking consent from patients is primarily a job responsibility of nurses in India.

Availability of adequate logistics is an essential part of pre-service trainings and helps students to adhere to the standard protocols with the clients. A similar study reported difficulties in delivering quality services as per the protocols, which were attributed to improper facility layout and lack of furniture [27]. Inadequate logistics and privacy may also have an effect on the client’s satisfaction [28, 29].

One of the main strength of our study was the use of reliable and valid method i.e., OSCE, to assess the skills of the study participants [30,31,32]. Some nurses refused to demonstrate the procedure of using a condom, which could be linked with shyness, or with unawareness of the correct procedure. This has implications on proper demonstration of condom usage to the clients, which may lead to higher failure rates and inconsistent use of condoms. This may also result in increased use of emergency contraceptives in the population at high risk of unintended pregnancy. Lim MS et al (China, 2015) has stressed that efforts should be made to sensitize the students to rise above and overcome this social taboo, and emphasis should be given to training in skills for condom/contraception negotiation, as partner refusal to use condoms is common [33]. The use of MEC wheel during pre-service training should be stressed upon as it aims to provide guidance to health-care providers in decision making and minimizing errors [34].

We did not assess the knowledge of the teachers regarding family planning services, which could have helped us to identify gaps in the medical education pattern; and of the participants regarding female condoms, which should be promoted to reduce the sexually transmitted infections [35]. Since this was a cross-sectional study, and information was self-reported hence, there could be a chance of recall and social desirability bias. Though interviews were conducted with one respondent at one time in closed room settings, yet response bias could be there due to prior interpersonal communication among the participants.

Based upon the results of this study it is recommended that training of interns and nurses on FP services should be given more emphasis during the medical education. Internship duties should include a minimum set of procedures and duties regarding FP services, and skills and knowledge acquired should be assessed by conducting the exit tests. Faculty members of the medical college should take the responsibility of teaching their students with a more serious approach towards FP services to ensure universal access to sexual and reproductive health care services and rights.

## Acknowledgments

We are thankful to Prof Rajesh Kumar, Dean (Academics), Professor and Head of Department of Community Medicine to provide guidance throughout the conduct of this study. We would like to extend our gratitude to Dr. Pravin H Shingare, Director of Medical Education and Research, Government of Maharashtra, Dr Ajay S Chandanwale, Dean, BJ Medical College Pune, Dr Nanandkar Sudhir Digambar, Dean, Grant Medical College and Sir JJ Group of Hospital Mumbai, Maharashtra; Dr S Ramji, Dean, Maulana Azad Medical College, New Delhi; Dr RK Ghokroo, Principal, Jawaharlal Nehru Medical College, Ajmer, Rajasthan, Dr SS Rathore, Dean, Sampuranand Medical College, Jodhpur, Rajasthan, for permitting us to conduct this study in their respective institutions within the study states. We would also like to thank Dr. Rajkrishna Ravikumar, for data collection in Rajasthan and Delhi.

## Additional Author information

### 1. Madhu Gupta, *MD, Ph.D*.^a^

^a^ Professor, Department of Community Medicine and School of Public Health Postgraduate Institute of Medical Education and Research, Chandigarh, 160012. India. Mobile: +917087008223; +917009769629.

### 2. Madhur Verma, *MD* ^a, b^

^a^ Senior Resident, Department of Community Medicine and School of Public Health, Postgraduate Institute of Medical Education and Research, Chandigarh, 160012

^b^ Assistant Professor, Department of Community Medicine, Kalpana Chawla Government Medical College, Karnal, Haryana. 132001. India. Mobile no:+919466445513.

### 3. Kiranjit Kaur, *MPH* ^a^

^a^ Research Officer, Department of Community Medicine and School of Public Health Postgraduate Institute of Medical Education and Research, Chandigarh, 160012. India. Mobile no:+918699291784.

### 4. Kirti Iyengar, *MD, PhD*^c^

^c^ National Programme Officer (Reproductive Health & HIV/ AIDs) United Nations Population Fund, 55 Lodi Estate, New Delhi 110003. India. Mob: +91 9799498350.

### 5. Tarundeep Singh, *MD* ^a^

^a^ Assistant Professor, Department of Community Medicine and School of Public Health Postgraduate Institute of Medical Education and Research, Chandigarh, 160012. India. **Email:**.

### 6. Anju Singh, *MD* ^d^

^d^ Assistant Professor, Department of Obstetrics and Gynaecology Postgraduate Institute of Medical Education and Research, Chandigarh, 160012. India.

## References

[i] Glasier A, Gülmezoglu AM. Putting sexual and reproductive health on the agenda. The Lancet. 2006 Nov 4;368(9547):1550–1.

[ii] World Population Prospects, The 2017 Revision: Data Booklet. Department of Economic and Social Affairs, Population Division (2017), United Nations. Available at http://www.un.org/en/development/desa/population/publications/databooklet/index.shtml.

[iii] The World Bank Data. India. Available from: https://data.worldbank.org/country/india. [cited: 2018, July 20].

[iv] Sample registration system. SRS Statistical Report 2016. New Delhi: Office of the Registrar General and Census Commissioner, Ministry of Home Affairs; 2016. Available from: http://www.censusindia.gov.in/vital_statistics/SRS_Reports_2016.html. [cited: 2018, July 20].

[v] International Institute of Population Sciences. Ministry of Health and Family Welfare. The government of India. National Family Health Survey. Round 4. India Fact Sheet. 2015-16. [cited 2018, May 14]. Available from: http://rchiips.org/NFHS/pdf/NFHS4/India.pdf.

[vi] Family Planning Division. Ministry of Health and Family Welfare Government of India. India’s ‘VISION FP 2020’. 2014. [cited 2018, May 14]Available from: https://advancefamilyplanning.org/sites/default/files/resources/FP2020-Vision-Document%20India.pdf.

[vii] Pachauri S. Expanding contraceptive choice in India: Issues and evidence. J Fam Welfare. 2004;50:13–25.

[viii] Ghooi RB, Deshpande S. Evidence-based medicine: A non-starter in India? Indian J Health Sci Biomed Res. 2016;9:121–6.

[ix] Kumar J, Hardee K. Rights-based family planning: 12 resources to guide programming. Resource guide. USAID. 2017. [cited 2018, May 14] Available from: http://ec2-54-210-230-186.compute-1.amazonaws.com/wp-content/uploads/2017/04/Resource-Guide-of-RBA-to-FP-Updated-April_2017.pdf.

[x] Dean AG, Sullivan KM, Soe MM. OpenEpi: Open Source Epidemiologic Statistics for Public Health, Version. www.OpenEpi.com, updated 2013/04/06, accessed 2018/07/05.

[11] Raselekoane NR, Morwe KG, Tshitangano T. University of Venda’s male students’ attitudes towards contraception and family planning. African journal of primary health care & family medicine. 2016;8(2):1–7.

[12] Kabagenyi A, Jennings L, Reid A, Nalwadda G, Ntozi J, Atuyambe L. Barriers to male involvement in contraceptive uptake and reproductive health services: a qualitative study of men and women’s perceptions in two rural districts in Uganda. Reproductive health. 2014 Dec; 11(1):21.

[13] Current family planning programme under public sector; National health mission. Ministry of Health and Family Welfare Government of India. 2018. [cited 2018, August 29]. Available from: http://nhm.gov.in/nrhm-components/rmnch-a/family-planning/background.html.

[14] Fehring RJ, Hanson L, Stanford JB. Nurse-midwives’ knowledge and promotion of lactational amenorrhea and other natural family-planning methods for child spacing. J Midwifery Women’s Heal. 2001;46(2):68–73.

[15] Jain AK. Information about methods received by contraceptive users in India. J Biosoc Sci [Internet]. 2017 Nov 8 [cited 2017 Dec 26];49(6):798–810. Available from: https://www.cambridge.org/core/product/identifier/S0021932016000602/type/journal_article

[16] Tilahun M, Mengistie B, Egata G, Reda AA. Health workers’ attitudes toward sexual and reproductive health services for unmarried adolescents in Ethiopia. Reprod Health. 2012;9(1):1–7.

[17] Nath A. HIV/AIDS and Indian youth—a review of the literature (1980–2008). SAHARA J 2009;6:2–8.

[18] Brown SS, Burdette L, Rodriguez P. Looking inward: provider-based barriers to contraception among teens and young adults. Contraception. 2008;78:355–7. 22.

[19] Campbell M, Sahin-Hodoglugil NN, Potts M. Barriers to fertility regulation: a review of the literature. Stud Fam Plann. 2006;37:87–98

[20] Najafi F. Investigation of Knowledge and Attitude of Family Health Workers about IUD, Norplant, and DMPA in Health Care Centres in East of Guilan Between 2001-2002. J Guilan Univ Med Sciences. 2004 15;13(50):14–21.

[21] van Zijl S, Morroni C, van der Spuy ZM. A survey to assess knowledge and acceptability of the intrauterine device in the Family Planning Services in Cape Town, South Africa. J Fam Plann Reprod Health Care. 2010;36(1):73–8.

[22] Rutter W, Knight C, Vizzard J, Mira M, Abraham S. Women’s attitudes to withdrawal bleeding and their knowledge and beliefs about the oral contraceptive pill. Med J Aust [Internet]. 1988 Oct 17 [cited 2018 Jan 9];149(8):417–9.

[23] Simbar M, Tehrani FR, Hashemi Z. Reproductive health knowledge, attitudes, and practices of Iranian. East Mediterr Heal J. 2005;11:888–97.

[24] Hogmark S, Klingberg-Allvin M, Gemzell-Danielsson K, Ohlsson H, Essén B. Medical students’ knowledge, attitudes and perceptions towards contraceptive use and counselling: a cross-sectional survey in Maharashtra, India. BMJ Open [Internet]. 2013. [cited 2018 Jan 1];3(12):e003739. Available from: http://www.ncbi.nlm.nih.gov/pubmed/24334156

[25] Hight-Laukaran V, Labbok MH, Peterson AE, Fletcher V, von Hertzen H, Van Look PF, et al. Multicenter Study of the Lactational Amenorrhea Method (LAM): II. Acceptability, Utility, and Policy Implications. [cited 2018 Jan 5]; Available from: http://irh.org/wp-content/uploads/2013/04/Hight-Laukaran_1997_multicenter_study_of_LAM_II.pdf.

[26] Singh S, Mittal S, Anandalakshmy PN, Goel V. Emergency contraception: Knowledge and views of doctors in Delhi. Heal Popul Perspect Issues. 2002;25(1):45–54.

[27] Atuahene MD, Afari EO, Adjuik M, Obed S. Health knowledge, attitudes, and practices of family planning service providers and clients in Akwapim North District of Ghana. Contraception and reproductive medicine. 2016;1(1):5.

[28] Ndulo J. Quality of care in sexually transmitted diseases in Zambia: patients’ perspectives. East Afr Med J. 1995;72(10):641–44. 32.

[29] Whittaker M, Mita R, Hossain B, Koenig M. Evaluating rural Bangladeshi women’s perspectives of quality in family planning services. Health Care Women Int. 1996;17(5):393–411.

[30] Grand’Maison P, Lescop J, Rainsberry P, Brailovsky CA. Large-scale use of an objective, structured clinical examination for licensing family physicians. Cmaj. 1992;146(10):1735–40.

[31] Bartfay WJ, Rombough R, Howse E, Leblanc R. Evaluation. The OSCE approach in nursing education. Can Nurse [Internet]. 2004 Mar [cited 2017 Dec 23];100(3):18–23. Available from: http://www.ncbi.nlm.nih.gov/pubmed/15077517

[32] Alinier G, Hunt B, Gordon R, Harwood C. Effectiveness of intermediate-fidelity simulation training technology in undergraduate nursing education. J Adv Nurs [Internet]. 2006. [cited 2018 Jan 1];54(3):359–69. Available from: http://doi.wiley.com/10.1111/j.1365-2648.2006.03810.

[33] Lim MS, Zhang XD, Kennedy E, Li Y, Yang Y, Li L, Li YX, Temmerman M, Luchters S. Sexual and reproductive health knowledge, contraception uptake, and factors associated with unmet need for modern contraception among adolescent female sex workers in China. PloS one. 2015 27;10(1):e0115435.

[34] World Health Organization. Medical Eligibility Criteria wheel for Contraceptive use. 2015. [cited 2018 Sep 21]. Available from http://www.who.int/reproductivehealth/publications/family_planning/mec-wheel-5th/en/.

[35] Mashanda-Tafaune B, Monareng LV. Perception and attitude of healthcare workers towards the use of a female condom in Gaborone, Botswana. health sa gesondheid. 2016 Dec 31;21:162–70.

